# Matrix production in *Bacillus subtilis* biofilms is localized to a propagating front

**DOI:** 10.1101/129825

**Authors:** Siddarth Srinivasan, Ioana D. Vladescu, Stephan A. Koehler, Xiaoling Wang, Madhav Mani, Shmuel M. Rubinstein

## Abstract

Bacterial biofilms are surface attached microbial communities encased in self-produced extracellular polymeric substances (EPS). Development of a mature biofilm must require coordinating cell differentiation and multicellular activity at scales much larger than the single microbial unit. Here we demonstrate that during development of *Bacillus subtilis* biofilms, EPS matrix production is localized to a front propagating at the periphery. We show that within the front, cells switch off matrix production and transition to sporulation after a set time delay of *∼*100 min. Correlation analyses of fluctuations in fluorescence reporter activity reveals that the front emerges from a pair of gene expression waves of matrix production and sporulation. The expression waves travel across cells that are immobilized in the biofilm matrix, in contrast to active cell migration or horizontal colony spreading. A single length scale and time scale couples the spatiotemporal propagation of both fronts throughout development, with the front displacement obeying a *t*^1/2^ scaling law. As a result, gene expression patterns within the advancing fronts collapse to self-similar expression profiles. Our results indicate that development of bacterial biofilms may be governed by universal wave-like dynamics localized to a self-similar front.

---

The vast majority of bacteria do not exist as solitary cells but within structurally complex communities known as bacterial biofilms [1]. From dental plaques to the alkaline hot springs of Yellowstone, bacteria in natural aquatic and terrestrial ecosystems [2–5] exist in these surface-associated aggregates encased in a self-produced matrix, known as the extracellular polymeric substance (EPS) [6–8]. The EPS matrix is primarily composed of exopolysachharides and proteins [9]. The production of EPS facilitates the construction of sophisticated threedimensional structures [10]. Additionally, the rigid scaffold supports the immobilization of individual microorganisms, allowing for cellular signaling and the creation of localized homeostatic zones [11]. Thus, the EPS matrix supports the biofilm’s robust physiology by facilitating reproducible spatial patterns of gene expression, cellular differentiation, and morphology in a manner analogous to multicellular organisms [12–15]. As a result, biofilms are capable of performing a plethora of sophisticated functions, including promoting surface adhesion and aggregation [16], enhancing mechanical rigidity [17], advanced architecture for water retention and uptake of nutrients [18], promoting cell-cell communication [19], and conferring enhanced antibiotic resistance [20]. Thus it is key for microorganisms to have a robust collective strategy for regulating biofilm matrix production and spore generation.

*Bacillus subtilis* is a convenient model bacteria to work with in view of the extensive single-cell molecular analyses focused on investigating the lineage of biofilm formation [21]. In the early stages of the *B. subtilis* biofilm developmental cycle, populations of cells initially expressing motility-related genes switch to synthesizing the EPS matrix [21, 22]. At later stages, a transition occurs from matrix production to the transcription of genes involved in sporulation, via a complex developmental process [23]. Detailed studies have investigated the physical aspects of *B. subtilis* biofilm growth, such as the biomechanics of colony expansion [24–26], formation of elevated fruiting bodies [27], and sustained oscillations in expanding colonies [28]. To attain a comprehensive description of the dynamics of colony growth, one must consider the variety of cell phenotypes within the biofilm. Thus, it is key to account the interplay between physical constraints and the spatial and temporal dynamic patterns of gene expression and cell-differentiation.

Here we show that taking into account the spatial and temporal distribution of the motile, matrix-producing and sporulating cell phenotype is critical to understanding biofilm development. Using a strain with fluorescent transcriptional reporters for these three cellular phenotypes, we find that following an early transient period, matrix production is largely restricted to an active zone at the periphery. The active zone propagates outwards radially, resulting in the materialization of a coherent localized front of matrix production near the leading edge of the growing colony. At its wake, a large subpopulation of cells turn off matrix production and start sporulating; thus, a second front of sporulation activity emerges behind the matrix front. Surprisingly, front propagation is not an outcome of active migration or spreading of matrix producing cells. Instead, these dynamics are a consequence of two traveling waves of gene expression. A first wave initiates matrix expression in cells near the leading edge of the biofilm. Resultantly, these matrix-producing cells are immobilized by the secreted biofilm matrix. The immobilized cells soon transition to a sporulation phenotype after a time delay of *∼*100 min. This occurs when the second wave, initiating sporulation, arrives. The process persists as the biofilm grows, guaranteeing uniform matrix production at the edge and giving rise to self-similar spatial patterns of gene expression. Over time, the front velocity decreases according to a simple *t*^-1/2^ scaling law.

## Significance Statement

Development of a mature bacterial biofilm requires coordinating the multicellular activity at scales much larger than the single microbial unit. Utilizing fluorescent reporters, we show that in *Bacillus subtilis* biofilms, the dynamics of matrix production and sporulation are restricted to the periphery, and are localized to a propagating front. We provide evidence that the dynamics of the biofilm’s growing front are self-similar and are a consequence of waves of gene expression in an immobilized field of cells, as opposed to being driven by cell migration. Thus, our results demonstrate that the biofilm’s collective state is determined at their periphery where the environmental influence is strongest.

## RESULTS AND DISCUSSION

### Measurement of gene expression

An important feature of *B. subtilis* biofilm development is that cell differentiation leads to several coexisting and spatially heterogeneous distributions of cell types within the biofilm [21, 29–31]. We restrict our focus to the dynamics of three important cell types: (i) motile cells that express the *hag* gene [32], (ii) EPS matrix producing cells that express the *tapA-sipW-tasA* operon [33], and (iii) sporulating cells that express the *sspB* gene [34]. We use a modified NCIB3610 *B. subtilis* strain (MTC871) [35] that harbors three transcriptional fusions of distinct fluorescent proteins with promoters of the aforementioned genes. Approximately 72 h after inoculation (see *SI Methods* for culturing conditions), our biofilms exhibit a characteristic spatial pattern of fluorescence in all three channels, as shown in Fig. 1A. Notably, the expression of the *tapA* gene is localized to a ring near the periphery of the biofilm, shown in Fig. 1A. Three-dimensional (3D) confocal imaging confirms the localization of matrix production near the periphery, as shown in Figs. 1B, and 1C. Taken together, our data suggest that while matrix is present throughout the biofilm, its production is largely localized to an annulus within *<* 2 mm from the leading edge of the biofilm. Additionally, 3D confocal imaging reveals a thin layer of residual matrix production activity near the agar interface, underneath the sporulating cells, not evident in our 2D measurements. In summary, while confocal imaging reveals additional features within the bulk of the biofilm, the dynamics of gene expression at the developing front are accurately captured by our 2D analysis (see Fig. 1D).

**Figure 1.**
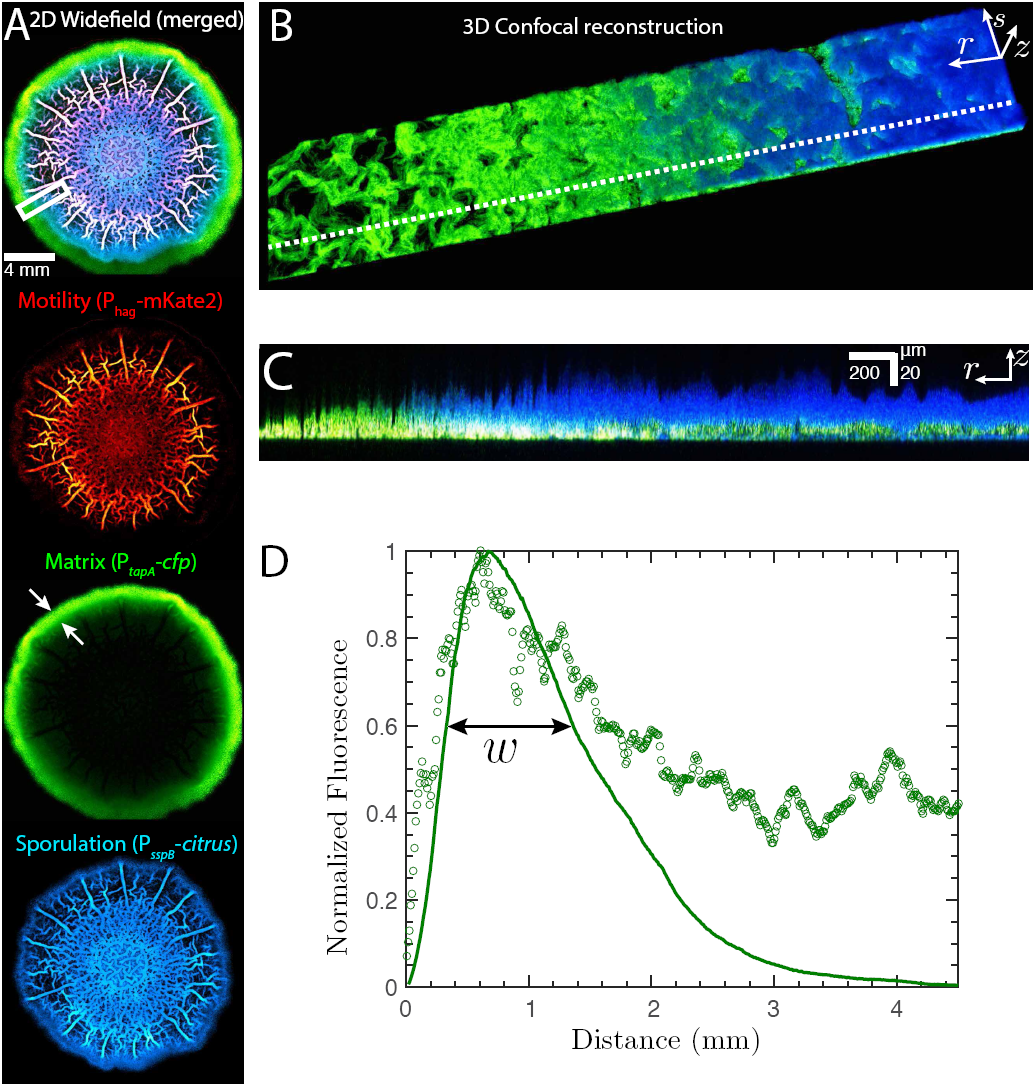
Comparison of fluorescent reporter expression profiles during *B. subtilis* colony expansion. (A) Superimposed 2D widefield image of a fluorescent triple-reporter *B. subtilis* biofilm colony at *t* = 72 h after inoculation. Individual background corrected fluorescent intensity maps correspond to the *P*_*hag*_*-mkate2* reporter of motility (red channel), the *P*_*tapA*_*cfp* reporter for matrix production (green channel), and the *P*_*sspB*_*-citrus* reporter for sporulation (blue channel). (B) 3D confocal reconstruction of a region of size *r × s × z* equals 4650 *µ*m × 470 × *µ*m 120 *µ*m of the same biofilm upon transferring to a coverslip. (C) Superimposed fluorescence activity of cells producing matrix (green) and sporulation (blue) along a single *rz* slice plane that corresponds to a cross-section of a zoomed out region along the dotted white line in Fig. 1B. (D) Comparison of the normalized fluorescence intensity profile as a function of distance from the biofilm edge. The solid curve corresponds to the 2D azimuthally averaged and normalized matrix channel profile from the widefield measurement. The data points represent normalized z-sum intensities averaged over 1280 separate *rz* slices. The width *w* is defined as the size of the region where the normalized fluorescence is greater than 60% of its maximum value.

### Matrix production localizes during biofilm development

As shown in Fig. 2A, during the first 24h after inoculation of biofilm development, the reporter for motility is localized to the central *∼* 1 mm zone of inoculation and is only weakly expressed outside (*<* 0.25 × peak expression, see Fig. 2B), whereas the reporter for matrix production is highly expressed throughout the entire colony. We note that these colony-scale observations are consistent with the switch from a motile to a sessile state observed in populations of single cells when exposed to biofilm inducing conditions [36, 37]. During the first 24h after inoculation, the reporter for sporulation is relatively weak in comparison to later stages of development (Fig. 2A).

**Figure 2.**
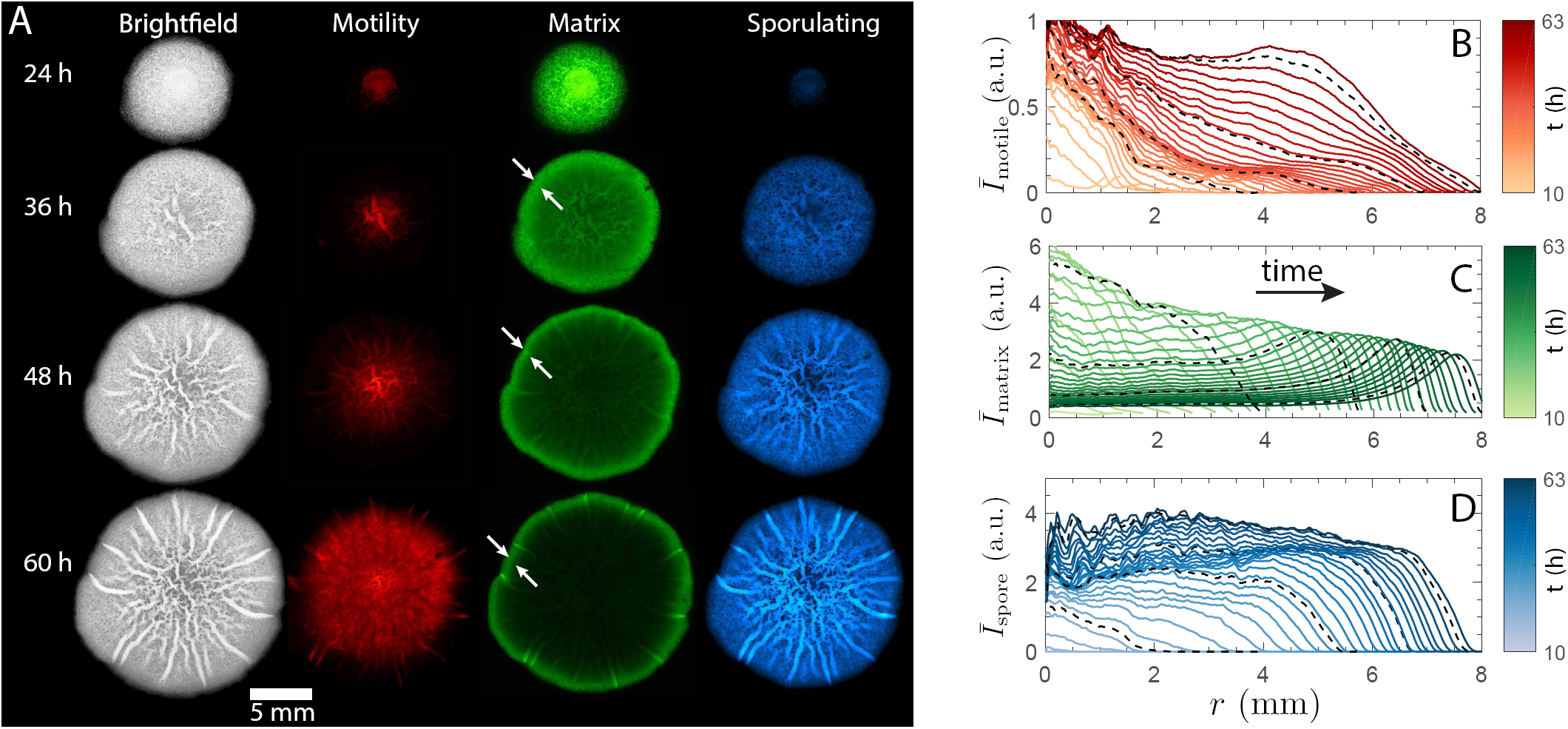
Timelapse of a developing biofilm and localization in EPS matrix production. (A) Background corrected fluorescent intensities 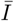 of the *P*_*hag*_*-mkate2* reporter of motility (red channel), the *P*_*tapA*_*-cfp* reporter for matrix production (green channel), the *PsspB-citrus* report for sporulation (blue channel) and the brightfield images, shown at intervals of 24h, 36h, 48h and 60h. (B-D) Azimuthally averaged profiles 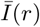 for the motility, matrix and sporulating channels, respectively. Averages are calculated over points equidistant from the center (see Fig. S1). Successive curves are 100 min apart, with darker curves corresponding to later times. Dashed lines correspond to the averaged profiles in each channel at 24 h, 36 h, 48 h and 60 h respectively.

As development proceeds (Movies S1-S4), matrix production steadily localizes to the periphery, as shown by the formation of a well-defined localized front of matrix reporter activity at 36 h in Fig. 2A. The emergence of the matrix front is a consequence of *tapA* expression levels falling off in the biofilm interior leading to a distinct peak of matrix expression near the periphery, as shown in Fig. 2C. The matrix front travels radially outwards. Behind the front, the expression of *tapA* drops by *>* 50%, while ahead of the front is uncolonized agar, as shown in Fig. 2C. Even during the formation of architecturally complex wrinkles, a morphological phenotype typically associated with robust biofilm formation [38], matrix production remains mainly localized to within a propagating front during expansion (Movie S2). As matrix production switches off in the interior, reporter activity levels of sporulation rises in an approximately uniform manner in space, as shown by Fig. 2D.

The onset of sporulation activity is closely associated with matrix production. The transition from matrix production to sporulation occurs predominantly in the vicinity of the front, as seen by the large gradients in 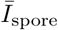 in Fig. 2D. At the latest observed stages of biofilm development, the reporter for motility rises dramatically within the biofilm interior, as seen for a 60 h colony in Fig. 2A, contrasting the repression of motility seen in single cells in biofilm inducing conditions [37]. Note that cells expressing the motility program within the biofilm interior are likely immobilized within the biofilm EPS matrix (Movie S5).

### Spatiotemporal dynamics of gene expression

Localization in *tapA* expression first appears as matrix production switches off within the interior of the biofilm. This defines spatially distinct, colony-scale, *‘on’* and *‘off’* regions, of extent 𝓁_*tapA*_(*t*), as shown in Fig. 3A. Similarly, the spatial extent of sporulation can be characterized by 𝓁_*sspB*_(*t*), and the extent of motility by 𝓁_*hag*_(*t*), as shown in Fig. 3A. Matrix production localizes to the region between 𝓁_*tapA*_(*t*) and *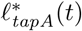* as indicated in Fig. 3A, and as observed in the confocal slice in Fig. 3B. Furthermore, sporulation activity follows matrix production as evidenced by the delayed onset of sporulation in Fig. 3B.

**Figure 3.**
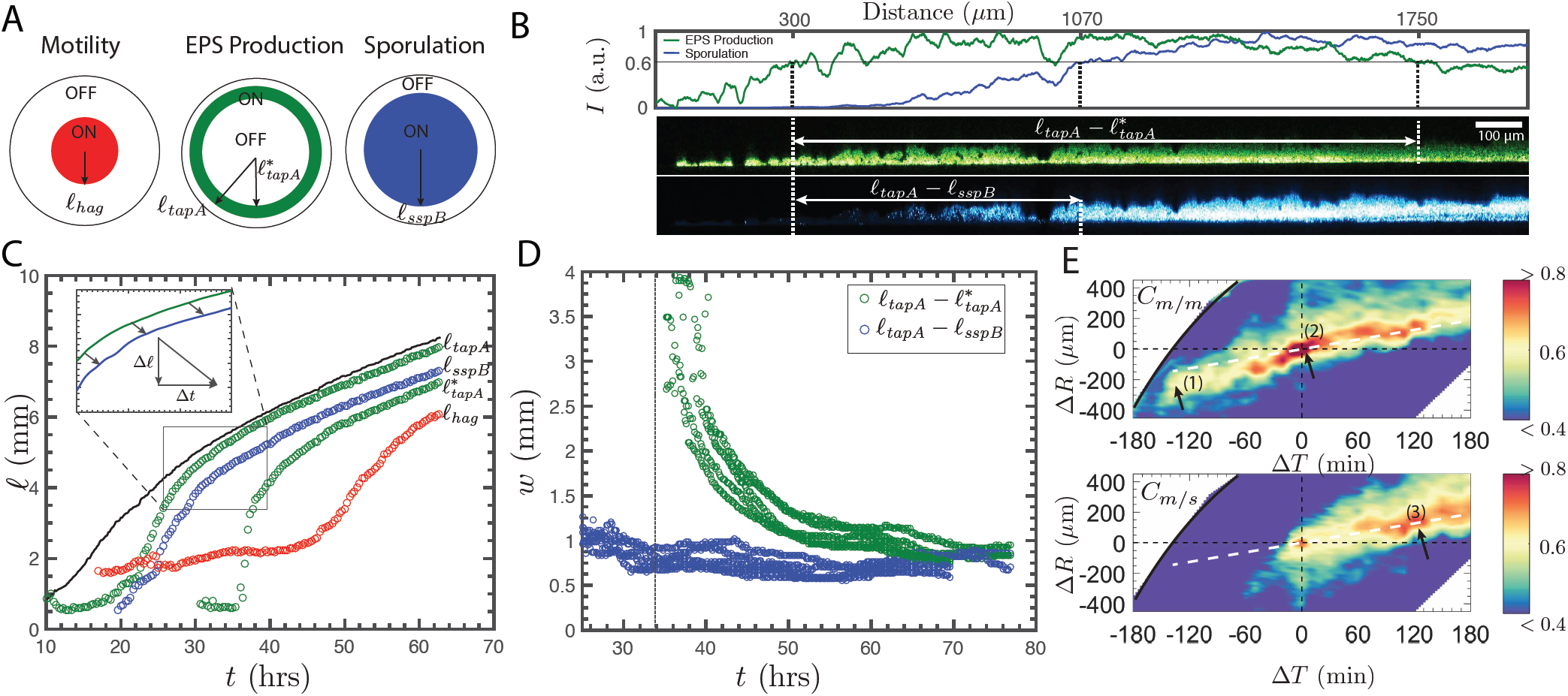
Spatiotemporal dynamics of the propagating front. (A) Schematic illustrating spatial localization of different-cell types within the biofilm. The spatial extents 𝓁_*hag*_(*t*) for motility, 𝓁_*tapA*_(*t*), *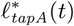* for matrix production, and 𝓁_*sspB*_(*t*) for sporulation are defined as distance at which *I*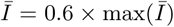 for each cell-type. For matrix production, 𝓁_*tapA*_(*t*) and *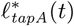* denote the inner and outer spatial bounds of localization. (B) Top: Normalized z-sum intensities from confocal measurements averaged of 420 *rz* slices in a 60 h biofilm. The vertical dashed lines indicate the location of 𝓁_*tapA*_, 𝓁_*sspB*_ and 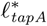. Bottom: Representative radial confocal slice showing the width of the matrix front, 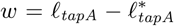, and the distance, 𝓁_*tapA*_*-𝓁*_*sspB*_, between the matrix and sporulation fronts. (C) Propagation of the spatial extents 𝓁_*tapA*_(*t*), 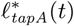 (green), 𝓁_*sspB*_(*t*) (blue) and 𝓁_*hag*_(*t*) (red) with time during development of a representative biofilm colony. The solid black line indicates the location of the biofilm edge. The two green curves represent the propagation of *RtapA* and *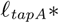* respectively. The inset shows complete overlap between 𝓁_*tapA*_(*t*) and 𝓁_*sspB*_(*t*) upon shifting by a length scale Δ 𝓁 = 550 *±* 100 *µ*m and a timescale Δ*t* = 1.4 *±* 0.5 h, calculated over *n* = 5 repeats. (D) Matrix front width *R*_*tapA*_ 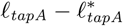 (in green) and the matrix-sporulation front distance 𝓁_*tapA*_ *𝓁*_*sspB*_ (in blue) for *n* = 5 repeats. (E) Matrix-matrix correlation coefficient *C*_*m/m*_ (top) and the matrix-spore correlation coefficient *C*_*m/s*_ (bottom) for a *T* = 24 h biofilm. Correlations are computed between fluctuations of reporter activity as detailed in *SI Methods* and Fig. S2. Positive values of Δ*R* indicate moving radially away from the matrix front towards the biofilm edge, and positive values of Δ*T* indicate later times. The solid black line denotes the biofilm edge. The dashed white line tracks the advection of clusters of cells within the biofilm during growth, as discussed in the main text.

The matrix production and sporulation fronts can be quantitatively characterized by a single length scale of Δ 𝓁 = 550 *µ*m*±*100 *µ*m and time scale of Δ*t* = 1.4 h*±*0.5 h (Fig. 3C). Specifically, 𝓁_*tapA*_(*t*) and 𝓁_*sspB*_(*t*) overlap when shifted by Δ 𝓁 and Δ*t* so that 𝓁_*tapA*_ = Δ 𝓁+ 𝓁_*sspB*_(*t-*Δ*t*), as shown in the inset of Fig. 3C. This suggests that the propagation velocities of the matrix production and sporulation fronts are well approximated 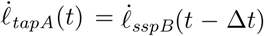 throughout colony expansion. The width of the matrix front 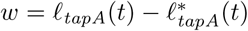 approaches a constant value of *∼* 1 mm. Similarly, the spatial distance, 𝓁_*tapA*_(*t*) *-𝓁*_*sspB*_(*t*), between the matrix production and sporulating fronts asymptotes to *∼*600 *µ*m, as shown in Fig. 3D.

### A traveling wave of gene expression

The dynamics underlying front propagation are revealed in the spatiotemporal correlations of fluctuations in reporter activity, as shown in Fig. 3E, and computed using the technique described in Fig. S2. The matrix production autocorrelation map, *C*_*m/m*_, as well as the matrix production sporulation cross correlation map, *C*_*m/s*_, is shown for clusters of cells that are initially located at the matrix production front *R* = 𝓁_*tapA*_(*T*) at *T* = 24 h. Correlation maps at later times are provided in Fig. S3. Positive values of Δ*R* correspond to regions away from the matrix production front and towards the edge, and positive value of Δ*T* indicate later times as measured in the stationary lab frame. Young cell clusters near the biofilm edge first began to produce matrix at Δ*T* = *-*120 min, as indicated in region 1 in Fig. 3E. Zones of high autocorrelation in the Δ*R -* Δ*T* plane track the propagation of these cell clusters (see *SI Methods* and Fig. S2). Subsequently, these cells are embedded within the biofilm EPS matrix and are only slowly advected outwards (relative to the biofilm edge) with a velocity 0.06 mm/hr, as indicated by the slope of the dashed white line in Fig. 3E. This advection velocity is also slower than the front propagation velocity 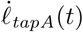, ensuing that the matrix production front overtakes the embedded cells at Δ*T* = 0 in region 2 in Fig. 3E. Meanwhile, the edge of the biofilm expands with a large velocity of 0.25 mm/hr at *T* = 24 h. Thus, the observed propagating matrix production front constitutes a wave in gene expression in a population of immobilized cells, contrasting cell migratory dynamics. Eventually, cells at the matrix production front switch to sporulation, as shown in region 3 in Fig. 3E. The transition to sporulation at Δ*T ∼* 100 min after peak matrix production in Fig. 3E is comparable to the 120 min time lag associated with the expression of the *sspB* gene following nutrient exhaustion in *B. subtilis* [39, 40]. This may indicate that the propagating waves of activity and the ensuing transitions we observe are driven by nutrient depletion.

### Front propagation demonstrates self-similar dynamics at late times

The asymptotic matrix production and sporulation profiles exhibit data collapse, indicative of universal dynamics, as shown in Fig. 4. These late stage universal matrix expression profiles are well-described by three features, a shift factor *R*^*\**^(*t*) that denotes the spatial location of the peak in matrix production, a scale factor *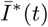* that measures the maximum production levels, and the shapes of the late stage asymptotic expression profiles. For a typical biofilm, the spatially shifted matrix production expression profiles steadily collapse onto a self-similar profile when translated by *R*^*\**^(*t*) and normalized *by 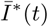,* as shown in Fig. 4A. Applying the same translation shift factor *R*^*\**^(*t*) to the sporulation channel also produces a data-collapse, as shown in Fig. 4B. Notably, the observed asymptotic profiles are quantitatively similar across different biofilms, as shown in Fig. 4C and Fig. 4D.

**Figure 4.**
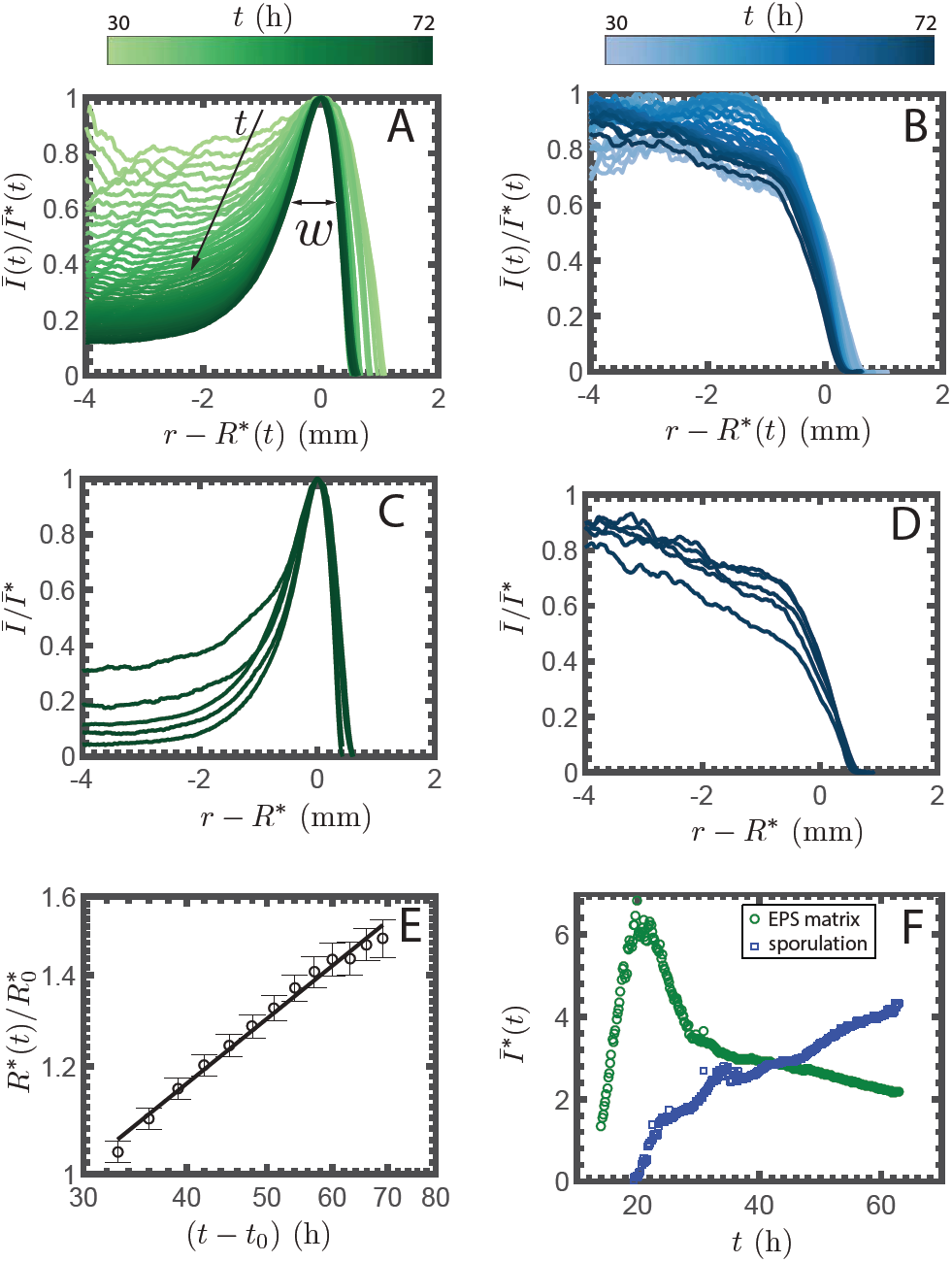
Self-similar profiles in a developing biofilm. (A) Spatial reporter activity profiles for matrix production, scaled by a factor *I*^*\**^(*t*) and shifted by a translation shift factor *R*^*\**^(*t*), that overlays the maximum intensities. *w* is the width of the matrix front. (B) Spatial profiles in the sporulation channel, where the same translation shift factor *R*^*\**^(*t*) collapses the curves. (C) Asymptotic late stage profiles (measured at *t >* 72 h) of matrix production for *n* = 5 independently grown biofilms. (D) Asymptotic late stage profiles of sporulation for *n* = 5 independently grown biofilms. (E) Translation shift factor *R*^*\**^(*t*) in logarithmic co-ordinates. The solid line indicates the best-fit scaling *R*^*\**^(*t*) *∼* (*t - t*0)^0^.^51^*±*^0^.^03^, where *t*0 = 10 h is the lag time before radial expansion. The shift factor is normalized by 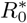, the initial onset position of the peak in matrix production. (F) Scale factors *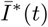* for matrix and sporulation reporter activity in a single representative biofilm.

The displacement of the propagating fronts scales as *R*^*\**^(*t*) *∼*(*t -t*_0_)^1/2^ as shown in Fig. 4E, where *t*_0_ = 10 h is the initial lag period prior to expansion. The normalization factor *I*^*\**^(*t*), shows a peak at *t*_*c*_ = 17 h, as shown in Fig. 4F. Beyond *t*_*c*_, global matrix production levels steadily drop within the biofilm, with a commensurate increase in sporulation. These results indicate that the patterns of gene expression at the periphery of a radially expanding biofilm are self-similar. In particular, the features highlighted here are observed regardless of differences in the initial inoculation event, and are thus universal. While we anticipate that varying growth conditions, and even strains, will quantitatively alter the observed self-similar dynamics, we might expect for the qualitative features (for example, the data collapse and the *t*^1/2^ scaling for the front displacement) to be preserved.

### The two regimes of radial expansion

The localization of matrix production to the propagating front leads to production of fresh biomass within this region. The physical expansion and spreading of the biofilm colony must result from mechanical forces associated with the production of biomass. We conclude with a quantitative characterization (see SI Methods) of two distinct regimes involved in radial biofilm expansion: an initial acceleration phase until *t*_*c*_, followed by a slowing-down phase beyond where the biofilm edge velocity *U*_max_ follows a well-defined scaling of the form *U*_max_ *∼*(*t - t*_*c*_)^-1/2^, as shown in Fig. 5. After the initial lag period *t*_0_ = 10 h, the biofilm expands radially. Colony expansion is very rapid at early times, as seen for a representative colony in Fig. 5A. For *t*_0_ *< t < t*_*c*_, the velocity progressively increases with distance from the biofilm interior to the exterior. The maximum colony expansion rate occurs at a critical time *t*_*c*_ = 17 h with a corresponding maximum velocity *U*_max_(*t*_*c*_) = 0.4 mm/h *±*0.04 mm/h. Note that *t*_*c*_ is identical to the global peak in matrix production (Fig. 4D). Beyond *t*_*c*_, there is a deceleration in expansion, indicated by the red profiles in Fig. 5A. Interestingly, despite both the biofilm edge velocity and the front velocity following a *t*^-1/2^ scaling at late times, the deceleration regime at *t*_*c*_ = 17 h occurs much earlier than the onset of localization in matrix production that is observed at 35 h *±* 2 h (Fig. 3C). This suggests that the dynamics which give rise to localization are distinct from the mechanics of expansion, the latter likely driven by biomass growth. Immediately behind the colony edge, we hypothesize that matrix production leads to aggregation of a thin layer of cells into 3-dimensional structures, that are slowly pushed out by the expanding biomass at the interior (Movie S6), and seen in the local dip behind the biofilm edge Fig. 5A. At late times the radial expansion is confined only to peripheral regions, as shown by the velocity magnitude map in the inset of Fig. 5A.

**Figure 5.**
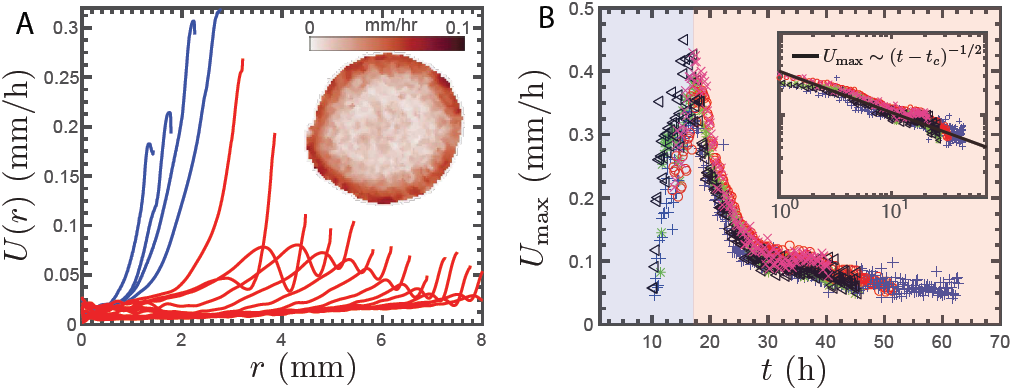
Dynamics of horizontal spreading during *B. subtilis* colony expansion. (A) Azimuthally averaged radial velocity profiles *U* (*r*). Successive curves are 100 min apart. Blue curves correspond to velocity profiles for *t < t*_*c*_ and the red curves for *t > t*_*c*_, where *t*_*c*_ = 17 *±* 2h is the characteristic time at which the maximum expansion velocity is attained. Inset: Plot of the 2D velocity magnitude map prior to azimuthal averaging, for a 36 h biofilm. (B) The evolution of *U*max as a function of time. The different colors indicate independently grown colonies under identical initial conditions. Inset: *U*max(*t*) in log-log. Solid line indicates *U*max *∼* (*t - t*_*c*_)^*-*^(^1/2^) where *t*_*c*_ = 17 h.

## Conclusions

We have demonstrated that matrix production and the onset of sporulation in a developing *B. subtilis* biofilm is not spatiotemporally uniform, but instead localizes to the cells within a propagating front at the biofilm periphery. The fronts are characterized by well-defined spatial patterns of matrix and sporulation reporter activities. Furthermore, our measurements reveals these propagating fronts are in fact waves of gene expression traveling via immobile bacteria, rather then active cell migration or horizontal colony spreading. Previously, it has been postulated that the spreading of biofilm colonies is a consequence of osmotic pressure generated by the production of the EPS matrix [24, 41, 42]. Such approaches assume uniform matrix production throughout the biofilm, and capture the initial vertical swelling and horizontal expansion, but do not allow for the observed spatial localization or predict the *t*^-1/2^ scaling of the expansion rate. The existence of a front suggests that colony expansion rate is nutrient-limited [43], and that the *∼* 1 mm width of the matrix-producing r<u>egio</u>n may be set by a nutrient penetration depth, 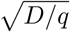, where *D* is the nutrient diffusivity and *q* is the nutrient uptake rate [44]. The accumulation of metabolic by-products, such as reactive oxygen species [45], has been recently shown to lead to decreased matrix gene expression in developing biofilms. Thus, metabolic gradients and nutrient-depletion dynamics likely drive the transition from matrix production to sporulation during front propagation.

It is widely believed that matrix production and biofilm formation confers enhanced fitness to the microbial community [46, 47]. While environmental variables such as metabolic gradients, oxidative stresses, osmotic pressure, fluid flow and antimicrobial agents are known to trigger specific genetic response circuits in individual bacterium [28, 48–50], the biofilm matrix enables the colony to adapt sophisticated response strategies at a colony scale. In natural settings, it is the matrix front at the biofilm periphery that directly interacts with changes in the local microenvironment. Therefore, our results may point to a broader strategy where collective decisions are made by the population at the front, and where sensitivity to changing environmental conditions is immediate.

## ACKNOWLEDGMENTS

We thank L. Mahadevan and M. Cabeen for discussions and suggestions. We thank M. Cabeen and R. Losick for providing strains. This work was supported by the National Science Foundation through the Harvard Materials Research Science and Engineering Center (DMR1420570). SMR acknowledges support from the Alfred P. Sloan research foundation.

## Supplementary Information: Supporting Information (SI)

### Bacterial strain

A modified NCIB 3610 *B. subtilis* strain, MTC871 was previously constructed from the wild-type RL2912 strain [35]. The MTC871 strain harbors three transcriptional fusion reporters 3610 *sacA::P*_*hag*_*-mkate2* [Kan^R^] *amyE::P*_*tapA*_*cfp*[Spc^R^] *ywrK::P*_*sspB*_*-citrus*[Cm^R^]. The mkate2 fluorescent protein reports on the activity of the promoter of the *hag* gene [32]. The cfp fluorescent protein reports on activity of the *tapA* gene [33]. The citrus fluorescent protein reports on the activity of the promoter of the *sspB* gene [34].

### Preparation of growth plates

All MTC871 biofilm colonies were inoculated and grown on 1.5 wt% agar gel plates infused with the MSgg biofilm promoting medium [27]. To prepare the agar plates, an intial 2x MSgg solution consisting of 6.15 mM potassium phosphate dibasic, 3.85 mM potassium phosphate monobasic, 200 mM MOPS buffer (pH 7), 4 mM MgCl_2_, 0.1 mM MnCl_2_, 2 *µ*M ZnCl_2_, 4 *µ*M thiamine, 100 *µ*g/ml phenylalanine, 100 *µ*g/ml tryptophan, 100 *µ*g/ml threonine, 1 wt% glycerol, 1 wt % glutamate, 1.4 mM CaCl_2_, 100 *µ*M FeCl_2_ was prepared and warmed to 55*°* C. A 3 wt% agar (A1296, Sigma) solution was separately prepared by autoclaving and cooled to 55*°* C. Equal volumes of the 2x MSgg solution and the 3 wt% agar solution were mixed and 50 *µ*g/ml Kanamycin was added at this stage. Finally, 6.7 ml of the 1.5 wt% agar/MSgg solution was poured into separate 35 mm petri dishes (*i.e.*, to obtain 7 mm thick agar plates) and cooled overnight at room temperature. Each 1.5 wt% agar/MSgg plate was dried for 10 mins in a laminar flow hood prior to use.

### Biofilm inoculation protocol

The MTC871 strain was incubated in fresh Luria-Bertani (Miller) medium in a 37*°* C incubator/shaker at 200 RPM for 4h. Subsequently, the culture was diluted to *OD*_650_=0.10. A 1 *µ*l drop of the diluted culture was deposited onto the center of a dried 1.5 wt% MSgg/Agar plate. The initial inoculation drop was then dried for 5 minutes and the colony was grown in a sealed petri dish to maintain a fixed relative humidity. After inoculation, the petri dish was transferred to the microscope stage for imaging. A custom built environmental chamber was used to maintain constant temperature during growth. A transparent Indium Tin Oxide resistive heater (HI-57Dp, Cell MicroControls) maintained at 30*°*C was used to heat the base and the lid of the petri dish to avoid condensation and allow for *insitu* imaging. A calibrated two-channel temperature control system (TC2BIP, Cell MicroControls), was used to provide 12V output to the resistive heater and maintain the set-point temperature to within *±*0.2*°* C.

### Widefield image acquisition and data analysis

As the biofilm colony matures, a sequence of widefield images is taken every 10 minutes over a period of 72 hours using a Zeiss Axiozoom.V16 microscope with a PlanNeoFluar Z 1.0x objective (NA 0.25), HXP 200 C metal halide illumination module, and a 16-bit Hamamatsu ORCA-Flash4.0 V2 Digital CMOS camera to detect the emitted light. The red mkate2 fluorescent protein was imaged using a Zeiss 63 HE filterset (Ex: BP 572/25; Em: BP 629/62), the cyan cfp fluorescent protein was imaged using a Zeiss 47 HE filter set (Ex: BP 436/25; Em: BP 480/40) and the yellow citrus fluorescent protein using a Zeiss 46 HE filter set (Ex: BP 500/25; Em: BP 535/30). There is negligible spectral overlap that allows for simultaneous imaging across all three channels as well as in brightfield. The algorithm to normalize the raw fluorescent images is shown schematically in Fig. S1. At every time point, each brightfield image is thresholded to obtain the foreground mask. Next, using the mask, the raw fluorescence images are normalized to obtain the background corrected intensity maps 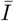, as shown in Fig. SI1 and in Fig. 2A in the main text. Finally, a Euclidiean distance map is obtained from the mask that specifies the distance of every pixel within the biofilm, *from the edge*. Finally, we azimuthally average the normalized intensities across all points that are equidistant from the center, as plotted in in Fig. S1. These averaged plots of 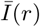 used to characterize the traveling fronts.

### Confocal imaging sample preparation

The mature biofilm growing on the agar gel is first gently peeled from the agar using a flat tip tweezer. We observed that the biofilm crumples like a thin elastic sheet during this process, but does not tear. Finally, we deposit the peeled biofilm onto a 50 mm MatTek glass bottom petri dish with a thin layer of DI water. When placed, the crumpled colony unfurls and floats on the thin water layer and regains its original morphology. We then remove the excess water using a pipette, until the biofilm colony is pressed flat against the glass bottom petri dish. Finally, we transfer the colony to confocal microscope for imaging. Confocal imaging was performed using a Leica SP5 microscope with a 63x water immersion objective (NA 1.2). A 458 nm argon laser-line was used as excitation for the cfp EPS reporter with a BP 485/15 detector channel, and a 543 nm HeNe laser-line was used as excitation for the citrus sporulation reporter with a BP 545/10 detector channel. The emitted light was recorded using an 8-bit PMT detector.

### PIV Analysis

The expansion of the biofilm was tracked using pairs of brightfield images acquired 10 mins apart with a 2048 pixel 2048 pixel field of view. For each pair of images, we used the MATLAB based PIVlab v1.4 app that implements an FFT cross-correlation based algorithm. We tracked displacements of a 64 64 pixel interrogation area using a 32-pixel step size. This corresponds to a 594 *µ*m x 594 *µ*m interrogation window with a 298 *µ*m step size. Extreme outliers in velocity, that correspond to noise/erroneous values, were rejected using a standard deviation and median filter in the post-processing validation step. Finally, the mean velocity magnitude was obtained by averaging across all pixels isodistant from the boundary (see Fig. S1D).

### Correlation matrix estimation

Correlations in the fluctuations of reporter activity are determined by partitioning the biofilm image into radial bins of 10 *µ*m, and angular bins of width Δ*θ* = 0.017 radians. Mean reporter activities, denoted as *I*^*i*^, are computed in each bin, as shown in Fig. S2. The subscript *i* = 1, 2*, … N* = (2*π/*Δ*θ*) corresponds to the angular position of each bin. At time *T*, the EPS front is located at *R* = 𝓁 _*tapA*_(*T*), as discussed in the main text. We define this as the reference location that corresponds to (Δ*R,* Δ*T*) = (0, 0). Therefore, at (0, 0), the vector 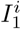 is calculated at the radial bin at the EPS front, as shown in Fig. S2B. Fluctuations in reporter activity are defined as 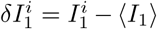, where the average is calculated over all angular locations. Expression patterns in the EPS and sporulation channel vary at time *T* + Δ*T*, as shown in Fig. S2A. Consequently, at a new spatiotemporal location (Δ*R,* Δ*T*) measured in the stationary lab frame, we determine mean reporter activity 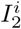 and fluctuations *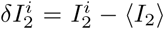* for the EPS channel (Fig. S2C) and the sporulation channel (Fig. S2D). From, *δI*_1_ and *δI*_2_, the spatiotemporal EPS-EPS auto correlation coefficients *C*_*m/m*_ and the EPS-sporulation cross correlation coefficients *C*_*m/s*_ are calculated as,

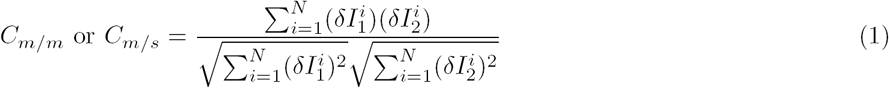

where *C*_*m/m*_ is the correlation between *δI*_1_ from the EPS channel initially located at the EPS front (0, 0), with *δI*_2_ from the EPS channel located at (Δ*R,* Δ*T*), as shown in Fig. S2C. Similarly, *C*_*m/s*_ measures correlations between *δI*_1_ from the EPS channel at (0, 0), with *δI*_2_ from the sporulation channel at (Δ*R,* Δ*T*), as shown in Fig. S2D.

**FIG. 1:**
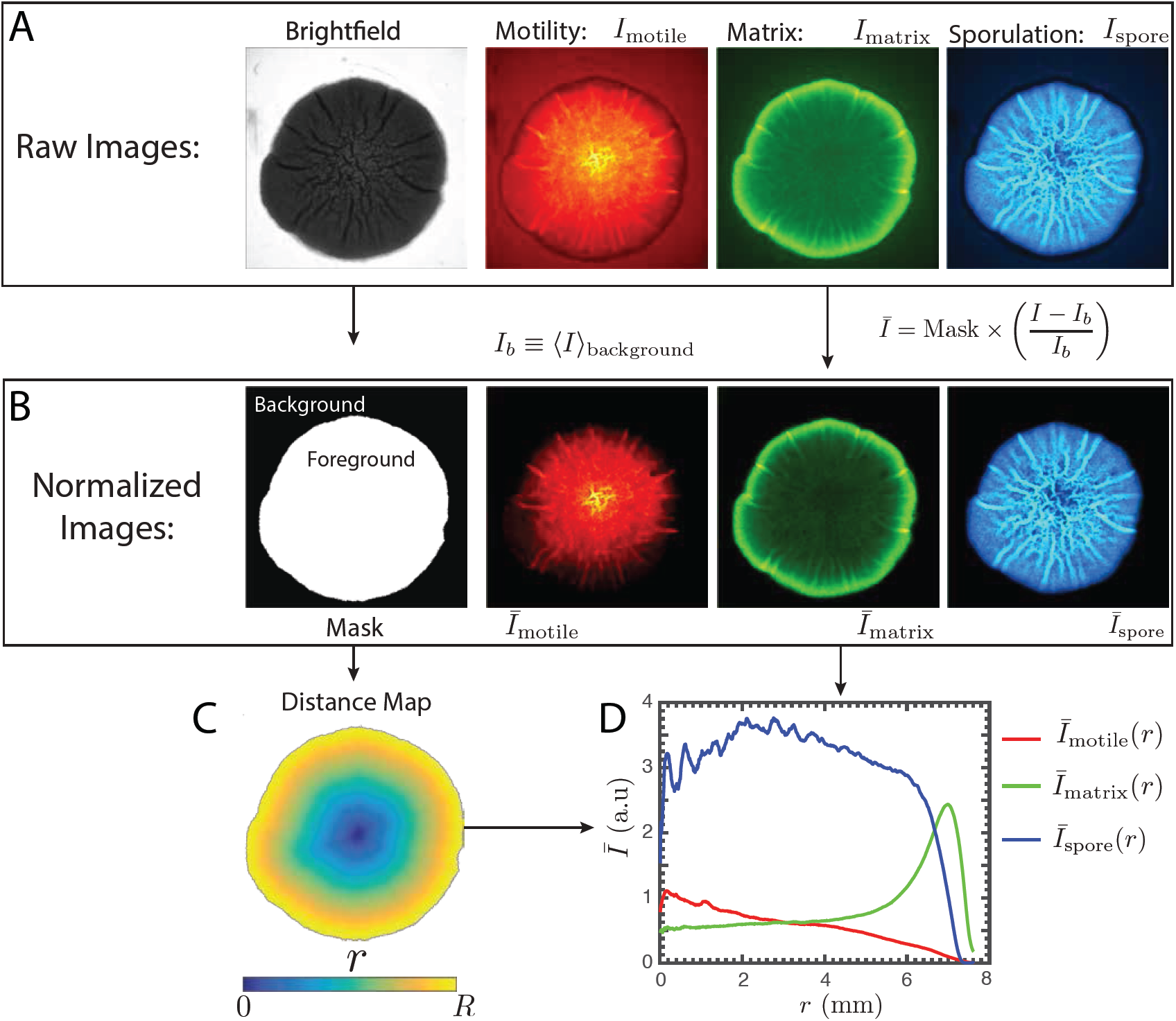
Normalized fluoresence activity and 1D reporter profiles. (A) Raw brightfield transmission and fluoresence images, *I*(*x, y*), corresponding to the *P*_*hag*_*-mkate2* reporter for motility (red channel), *P*_*tapA*_*-cfp* reporter for EPS production (green channel) and the *PsspB-citrus* reporter for sporulation (blue channel). (B) (Left): Image of the binary mask generated from the brightfield transmission image by Otsu thresholding. (Right) Images of the background subtracted and normalized foreground fluorescence images, 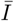, that are obtained from the raw images by the operation 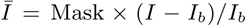. Here *I*_*b*_ is the mean background intensity that is obtained from averaging the signal in the background region for each channel. (C) Euclidean distance map generated from the binary mask. The colormap denotes the radial distance of all pixels that are equidistant from the edge. (D) Azimuthally averaged 1-D radial profiles corresponding to the motility (red), EPS production (green) and sporulation (blue) reporter activities. The averages are performed over all points equidistant from the edge using the Euclidean distance map to preserve symmetry.

**FIG. 2:**
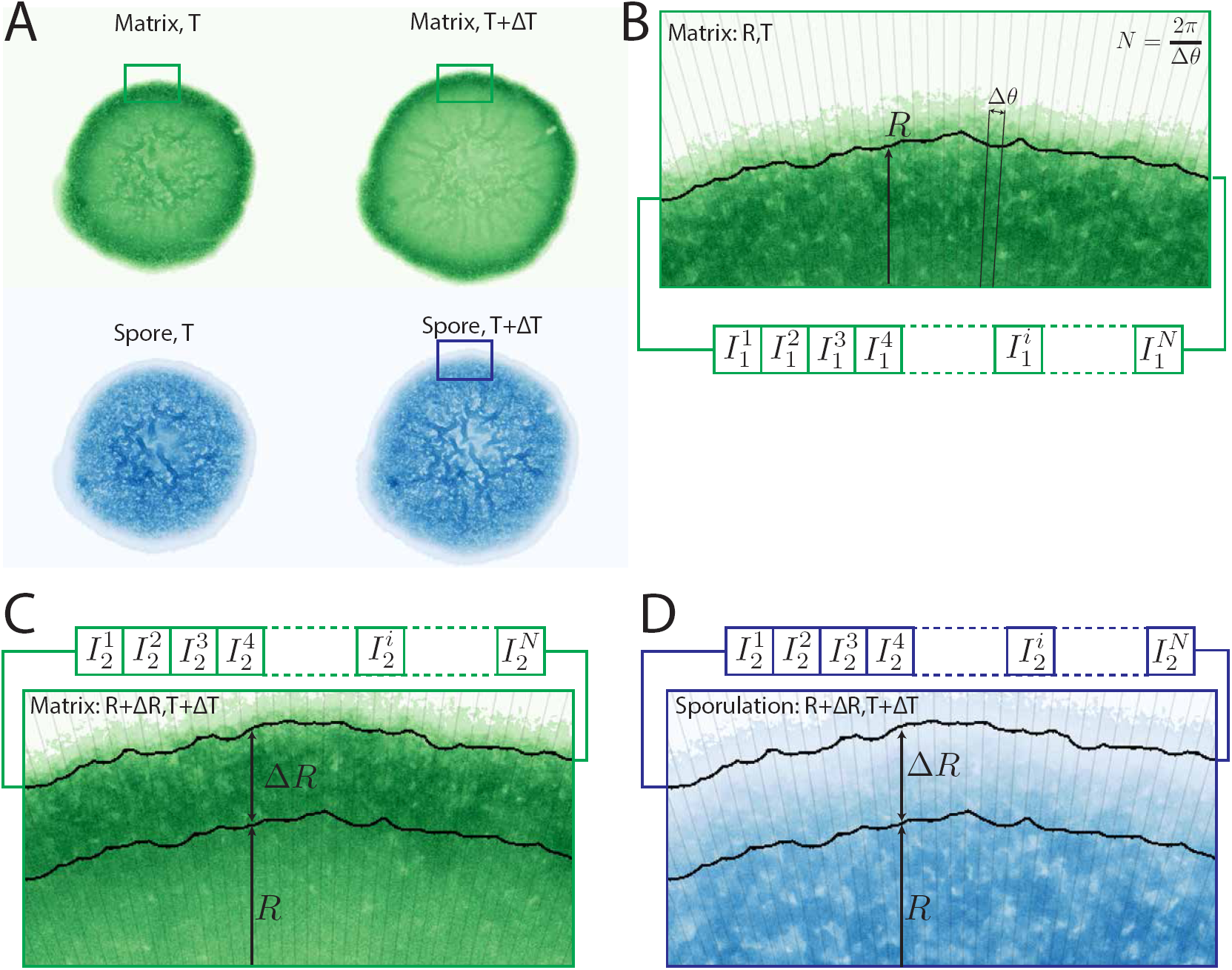
Determination of reporter activity fluctuations. (A) Images of the raw fluoresence intensities, *I*, corresponding to the EPS production and sporulation channels for a biofilm shown at at time *T* (left), and *T* + Δ*T* (right). (B) Region near the periphery at time *T* in the EPS channel. Mean intensities are estimated in bins of radial width 10 *µ*m and angular width Δ*θ* = 0.017 radians. The vector 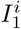 corresponds to the mean intensities at the radial bin located at the EPS front located at *R* = 𝓁*tapA*(*T*), as indicated by the solid black line. The subscript denotes the angular bin position. The fluctuations in reporter intensity are 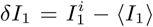, where the average is carried out over all azimuthal bins. (C) Region near the periphery of the biofilm at time *T* + Δ*T* in the EPS channel. The vector *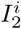* corresponds to the mean intensities in the EPS channel at a radial bin located at a new spatial location *R* + Δ*R* in the stationary lab frame. (D) Region near the periphery of the biofilm at time *T* + Δ*T* in the sporulation channel. Here, the vector 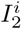 corresponds to the mean intensities in the sporulation channel at *R* + Δ*R* in the stationary lab frame.

**FIG. 3:**
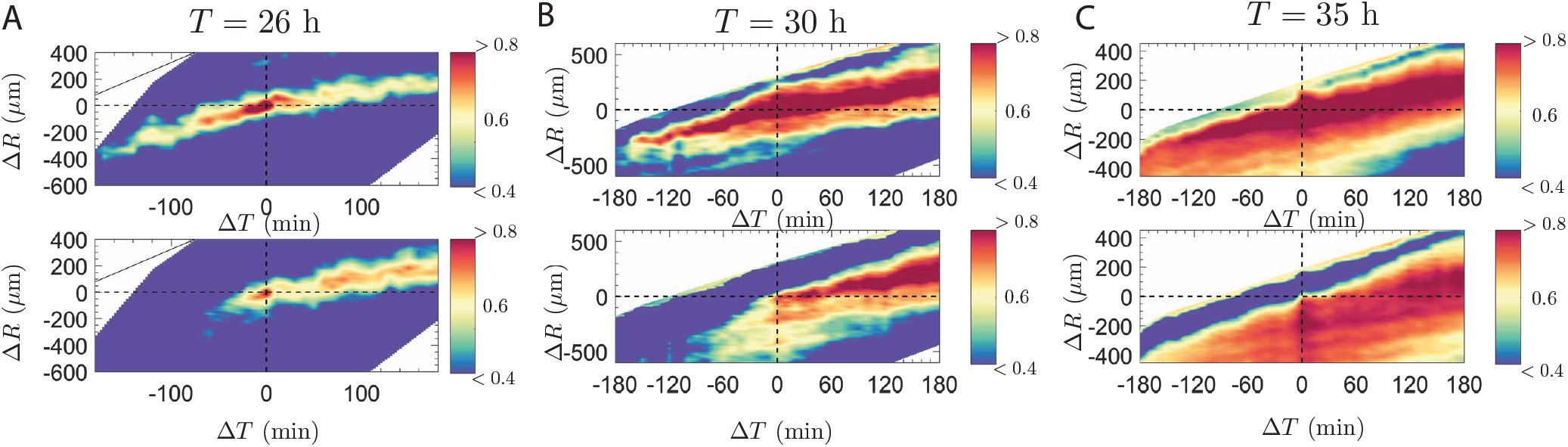
Correlation maps during biofilm development. Plots of the EPS-EPS correlation maps *C*_*m/m*_(Δ*R,* Δ*T*) (top) and the EPS-sporulation correlation maps *C*_*m/s*_(Δ*R,* Δ*T*) (bottom) at (A) T=26 h, (B) T=30 h and (C)T=35 h during colony expansion. *C*_*m/m*_ and *C*_*m/s*_ are calculated from the reporter fluctuations using equation (1). Zones of red track regions of high correlation during colony expansion. The EPS front initiates localization at *T* = 30, and is fully localized by *T* = 35 as discussed in the main text.

### SI Movies

**Supplmental Movie 1.** Movie of development of *B. subtilis* biofilm in the brightfield transmission channel at intervals of 10 mins. The movie shows the logarithm of the transmitted intensity using an inverted grayscale colormap.

**Supplmental Movie 2.** Movie of development of *B. subtilis* biofilm in the *P*_*tapA*_*-cfp* reporter channel for EPS production at intervals of 10 mins.

**Supplmental Movie 3.** Movie of development of *B. subtilis* biofilm in the *P*_*sspB*_*-citrus* reporter channel for sporulation at intervals of 10 mins.

**Supplmental Movie 4.** Movie of development of *B. subtilis* biofilm in the *P*_*hag*_*-mkate2* reporter channel for motility at intervals of 10 mins.

**Supplmental Movie 5.** Video of overlay of motile cells (red) and matrix producing cells (green) at the biofilm periphery at intervals of 100 msec. Cells expressing the motility genes are often jammed within the biofilm matrix.

**Supplmental Movie 6.** Zoom in movie of colony expansion at the edge of the biofilm showing the transition from sliding at the colony edge, to aggregation into 3-dimensional structures.

